# Aberrant tissue stiffness impairs neural tube development in Mthfd1l mutant mouse embryos

**DOI:** 10.1101/2023.08.04.552024

**Authors:** Yogeshwari S. Ambekar, Carlo Donato Caiaffa, Bogdan Wlodarczyk, Manmohan Singh, Alexander W. Schill, John Steele, Salavat R. Aglyamov, Giuliano Scarcelli, Richard H. Finnell, Kirill V. Larin

## Abstract

Neurulation is a highly synchronized biomechanical process leading to the formation of the brain and spinal cord, and its failure leads to neural tube defects (NTDs). Although we are rapidly learning the genetic mechanisms underlying NTDs, the biomechanical aspects are largely unknown. To understand the correlation between NTDs and tissue stiffness during neural tube closure (NTC), we imaged an NTD murine model using optical coherence tomography (OCT), Brillouin microscopy, and confocal fluorescence microscopy. Here, we associate structural information from OCT with local stiffness from the Brillouin signal of embryos undergoing neurulation. The stiffness of neuroepithelial tissues in Mthfd1l null embryos was significantly lower compared to that of wild-type embryos, while exogenous formate supplementation improved tissue stiffness and gross embryonic morphology in both nullizygous and heterozygous embryos. Our results demonstrate the significance of proper tissue stiffness for normal NTC and pave the way for future studies on the mechanobiology of normal and abnormal embryonic development.

## Introduction

Folate (vitamin B9) is an essential nutrient to support healthy cellular development. Folate-dependent one-carbon metabolism is involved in several core biological processes ranging from genome replication and cell division and is required to produce universal methyl donors, regeneration of redox cofactors, *de novo* purine and thymidylate synthesis, DNA synthesis, and amino acid metabolism(*1, 2*). During the first few weeks of pregnancy, the embryonic stem cells on the blastocyst’s inner cell mass extensively divide and differentiate into a variety of cell types and tissues to form the axial body plans of an early embryo. At this stage of embryonic development, folate is a critical requirement as an essential nutrient in the maternal diet(*3-5*). The human body does not produce folate on its own, and to sustain natural cell development and metabolic functions, folate must be obtained from nutrition (NIH Fact Sheet, 2022; WHO Executive Board, 2023). Folate deficiency during pregnancy is a major factor leading to the incidence of neural tube defects (NTDs), and adequate intake of folate during the periconceptional period can reduce the prevalence of this birth defect(*5*).

NTDs occur when the neural plate, an ectoderm-derived structure growing adjacent to the notochord and prechordal mesoderm, fails to fold and shape the neural tube at specific dorsally-located neuropores along its antero-posterior axis resulting in abnormal rostro-caudal development of the brain and the spinal cord (Figure 1). This can result in conditions such as spina bifida occulta (abnormal closure of the caudal neuropore with no herniation), spina bifida cystica (caudal neuropore fails to close, developing herniation at meninges and neural tissue), anencephaly (abnormal closure of the rostral neuropore leading to brain and skull defects), several brain abnormalities, outflow tract cardiac defects, and cleft lip or cleft palate(*1, 4, 6-9*). Folate is a critical component of one-carbon metabolism, facilitating the cycling of one-carbon units in the form of methyl groups, formyl groups, formaldehyde, or formate, and their production is regulated by numerous enzymes and cofactors, including vitamins B6, B9, and B12. One-carbon metabolism is linked to the folate and methionine cycles and is compartmentalized between the cytoplasm, mitochondria, and nucleus(*10*). Generally, carbon units are metabolized from donors such as serine or glycine in the mitochondria and are enzymatically oxidized to formate in a folate-mediated process. The mitochondrial-produced formate is then utilized in the cytoplasm or nucleus for the synthesis of purines, pyrimidines, and methionine. The methionine cycle occurs primarily in the mitochondria to generate and recycle methionine. In the nucleus, the one-carbon units are used as a source for DNA methylation, which is essential for epigenetic inheritance and regulation of gene expression. Impairment of one-carbon metabolism can lead to a variety of health problems, including NTDs, cardiovascular disease, and certain forms of cancer(*11*).

**Figure 1.**
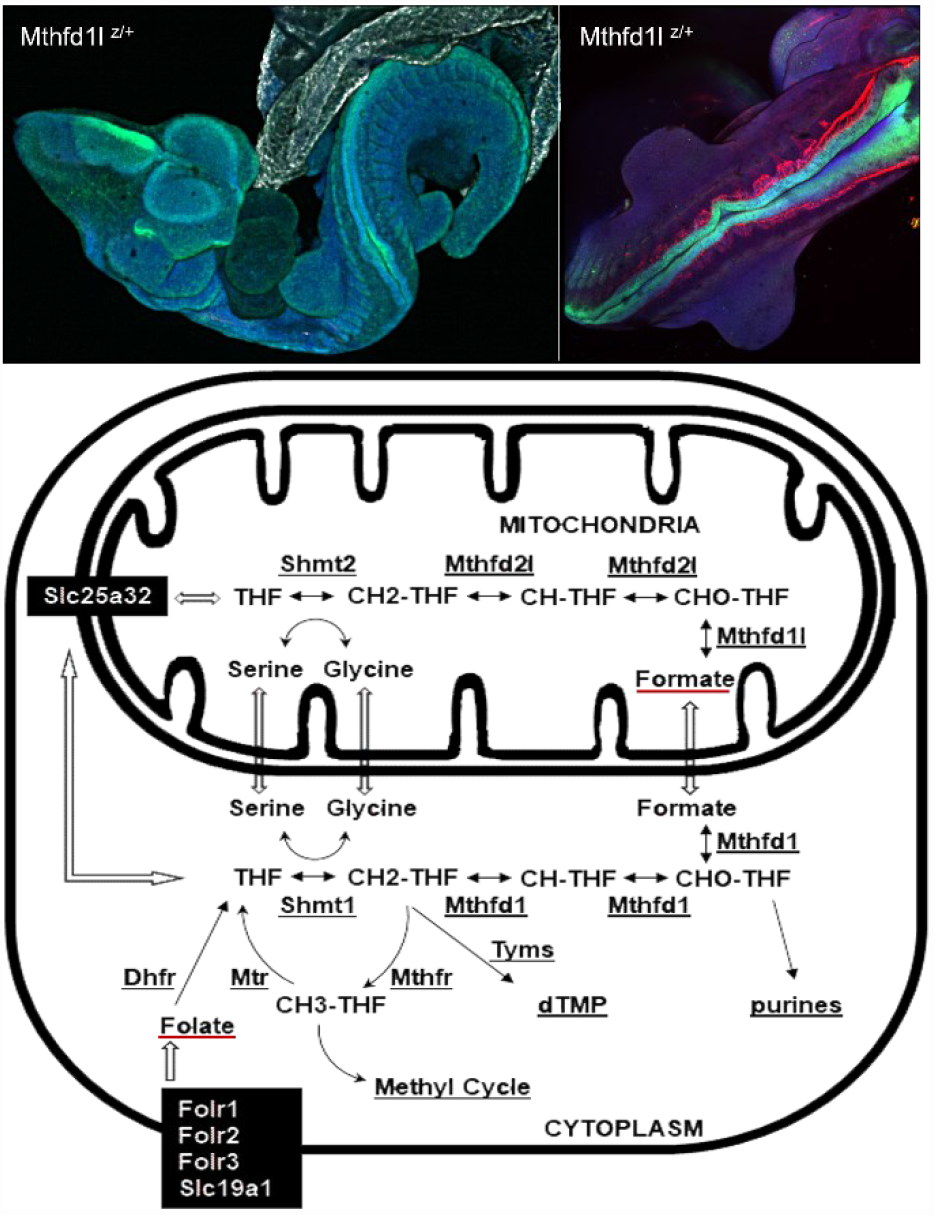
The One Carbon metabolism pathway is required for neurulation. Neural tube defects are a highly penetrant phenotype in the Mthfd1l knockout mice lineage. Failures during NTC are identified by immunostaining against Pax6, in green at the top images, which is highly localized to the neural tube. The diagram below represents the compartmentalization of the one-carbon metabolism in between the cytosol and mitochondria. The gene *Mthfd1l* encodes a mitochondrial monofunctional enzyme responsible for catalyzing 10-formyl-tetrahydrofolate to formate, which is the last step in the flow of one-carbon units from the mitochondria to the cytoplasm.

Methylenetetrahydrofolate dehydrogenase (NADP+ dependent) 1-like (*Mthfd1l*) is a gene encoding a mitochondrial monofunctional enzyme responsible for catalyzing the interconversion of 10-formyl-tetrahydrofolate to formate, which is the last step in the flow of one-carbon units from the mitochondria to the cytoplasm (Figure 1). Embryos lacking *Mthfd1l* have a significantly reduced capacity for mitochondrial formate production(*12*). *Mthfd1l* is expressed during all stages of mouse embryonic development, presenting higher levels of expression along the neural tube antero-posterior axis, adjacent paraxial mesoderm, craniofacial tissues, brain, limbs, and tail buds from embryonic (E) days 9.5 to 13.5(*13, 14*). Genetic variants or alterations in the expression levels of *Mthfd1l* in humans have been associated with cardiovascular disease, neurological conditions, cancer, and NTDs(*15-31*).

In this context, we focused on studying the effect of mutations of gene *Mthfd1l*. Deletion of *Mthfd1l* impairs normal organogenesis during embryonic development; nullizygous mouse embryos exhibit a characteristic growth delay, aberrant NTC, abnormal craniofacial development, and embryonic lethality by E12.5. The NTD phenotype is variable, including defects such as craniorachischisis, exencephaly, and a wavy neural tube(*14*). In addition, Shin and colleagues showed that E8.5 *Mthfd1l* mutant embryos present decreased cellular density at the head mesenchyme, in an early organogenesis stage where the first point of closure of the hindbrain neuropore is being formed(*32*). At this stage, apoptosis and neural crest cell specification were not affected by *Mthfd1l* ablation. Supplementation of pregnant dams with formate rescued mesenchymal density and cell proliferation in the nullizygous embryos(*32*). These findings suggest that NTDs resulting from impairment of one-carbon metabolism may be caused by alterations in local tissue stiffness disrupting the biomechanical properties that contribute to NTC.

While our understanding of the genetic and molecular mechanisms of neural tube closure (NTC) and NTDs has rapidly grown in recent years, the biomechanical processes leading to proper or aberrant NTC remain largely unknown. Yet, neurulation involves dramatic morphological changes that are widely accepted to be the result of a delicate balance between mechanical forces and tissue stiffness(*33*), i.e., the resistance to deformation under an applied force. However, it has been extremely challenging to map the biomechanical properties of tissues with high resolution *in vivo* using a noninvasive method during embryonic development.

In this study, we demonstrate a novel, multimodal imaging technique combining optical coherence tomography (OCT) and Brillouin light scattering microscopy to correlate neural tube structural and biomechanical changes in a murine NTD model. Brillouin microscopy is a noninvasive optical imaging technique capable of mapping the Brillouin frequency shift of tissues with high spatial resolution without contact(*34-36*). Imaging structural information is crucial for understanding the physical location of the captured Brillouin frequency shift to correlate embryo morphology with the mechanical information provided by the Brillouin frequency shift. OCT is a well-established technique that can noninvasively provide 3D structural details of developing embryos with high resolution(*37-39*). Earlier studies have demonstrated the feasibility of biomechanical assessment of the developing neural tube in mouse embryos using Brillouin microscopy(*40-42*). However, Brillouin microscopy does not provide structural information, which often results in lengthy alignment and imaging times, which are not suitable for *in vivo* studies. The inclusion of structural guidance would significantly speed up the imaging time and repeatability of Brillouin microscopy measurements (e.g., imaging the same region in multiple samples). Hence, we have developed a co-aligned Brillouin-OCT system with customized instrumentation software(*42*). Here, we have adapted, optimized, and combined two technologies to build a single Brillouin-OCT instrument for structurally guided tissue stiffness mapping of the neurulation process in *Mthfd1l* knockout mouse embryos with high resolution.

We demonstrate here that tissue stiffness is significantly lower in the neural tube neuroepithelia, at the otic pit, non-neural surface ectoderm, and adjacent paraxial mesenchyme of *Mthfd1l* mutants in comparison to similar-stage wild-type embryos. The neuronal fate of Pax6-positive neural progenitors was disrupted in these embryos, indicating that mitochondrial formate is not only important to sustain the contractile properties of the cytoskeleton and general extracellular matrix composition but also has a critical role during mammalian cell differentiation. The importance of mitochondrial formate during cell differentiation was also revealed by Pax3 staining, indicating that somitogenesis is disrupted in *Mthfd1l* mutants. Our data also indicate decreased migration of Sox10 positive neural crest cells along the embryo antero-posterior axis in the *Mthfd1l* mutants. Furthermore, we show that maternal supplementation with formate rescues tissue stiffness at the neural tube neuroepithelia, otic pit, non-neural surface ectoderm, and adjacent paraxial mesenchyme and reestablishes the potential of neural progenitor cell differentiation, somitogenesis, and neural crest migration in the supplemented *Mthfd1l* knockout embryos.

## Results

### Brillouin-OCT imaging during neural tube development in *Mthfd1l* mutants

Ablation of the *Mthfd1l* gene in mice disrupts normal neural tube development(*14, 32*). Due to a variety of aberrant neural tube phenotypes observed in embryos of this mouse line, we hypothesized that tissue stiffness is an intrinsic component driving failed NTC. Therefore, we took advantage of a novel multimodal imaging technique, relying on the association of high-resolution OCT and Brillouin microscopy to measure stiffness during anterior NTC. Three-dimensional OCT images were acquired to locate a region of interest on the embryonic neural tube with 1000 A-lines per B-scans, 1000 B-scans per volume, and 5 frames per position for averaging and signal-to-noise ratio (SNR) improvement. In addition to structural imaging and morphological phenotyping of the embryos, the OCT images were used to ensure a comparable region was imaged for all embryos. Two-dimensional OCT images consisting of 1000 A-lines per B-scan were also acquired to guide Brillouin imaging at a transverse plane throughout the otic pits in murine embryos at the developmental stages E9.5 (Figure 2, h-n) and E10.5 (Figure 3 g-l). Using the 2D-OCT images as a starting point, a transverse section crossing the otic pits was chosen to capture Brillouin light scattering in terms of Brillouin frequency shift. The dimensions of the Brillouin scans varied based on the size of cross-section selected using the 2D-OCT image on each embryo. The dimensions of the 2D Brillouin scan for wild-type and heterozygous embryos were ∼1 mm x ∼0.6 mm, and for nullizygous embryos, it was ∼0.5 mm x ∼0.5 mm, and Brillouin images were acquired in the selected region. The average Brillouin frequency shift was calculated based on the identification of tissues using 2D-OCT images (Supplementary Fig. 1).

**Figure 2.**
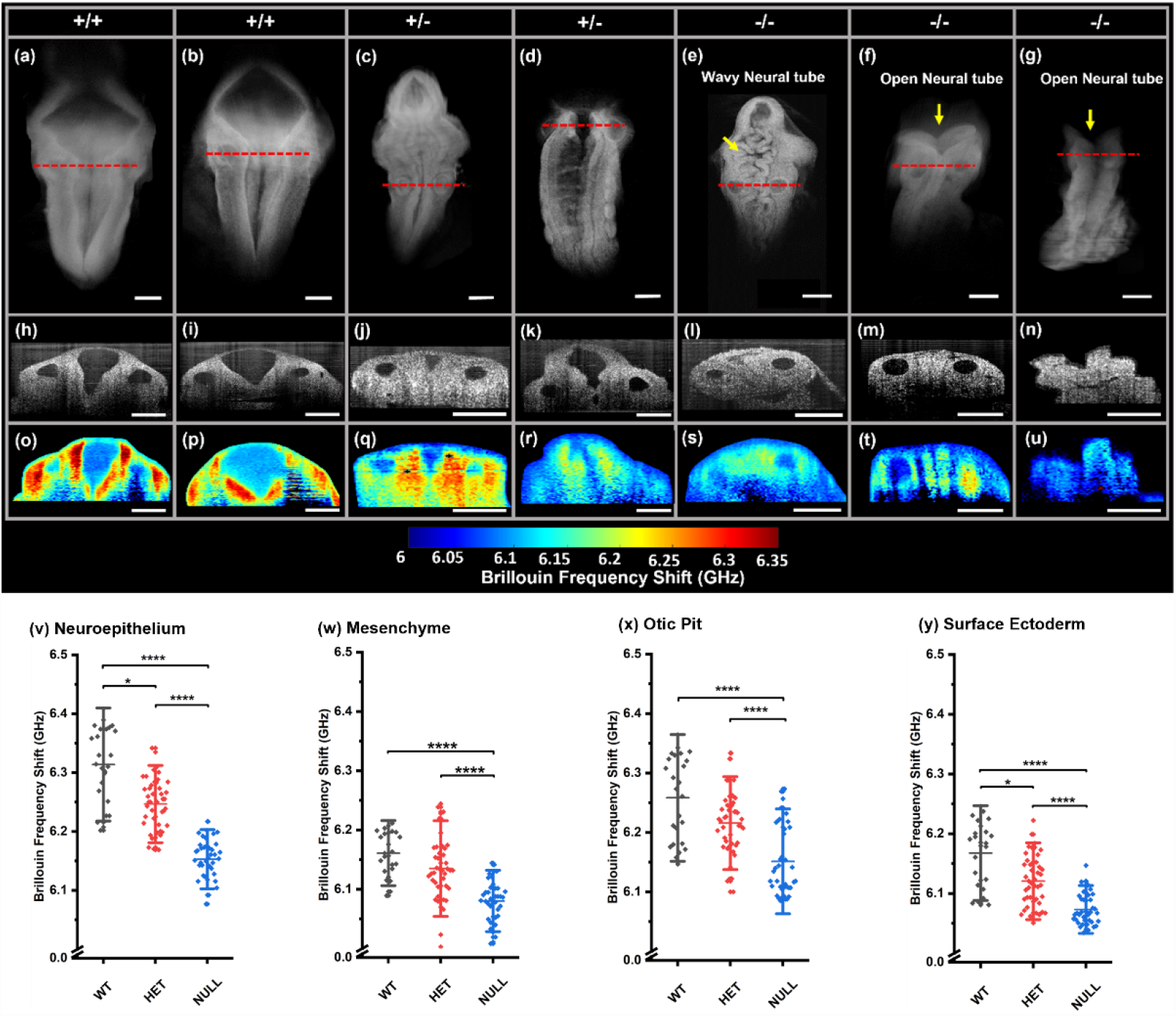
Mthfd1l ablation decreases tissue stiffness in E9.5 embryos. (a-g) 3D-OCT images showing the hindbrains of Mthfd1l embryos at E9.5. (h-n) 2D-OCT optical sections showing the neural folds at rhombomere 5 located near the otic pits. (o-u) Brillouin frequency shift images represent tissue stiffness in the anatomical areas identified by the corresponding 2D-OCT optical sections. Region-wise average Brillouin frequency shift of wild-type (WT), heterozygous (HET), and nullizygous (NULL) embryos at the (v) neural tube neuroepithelia, (w) adjacent paraxial mesenchyme, (x) otic pit, and (y) non-neural surface ectoderm region. * and **** indicate P < 0.05 and P <0.0001, respectively by pair-wise Dunn’s test. Scale bar is 0.25 mm.

**Figure 3.**
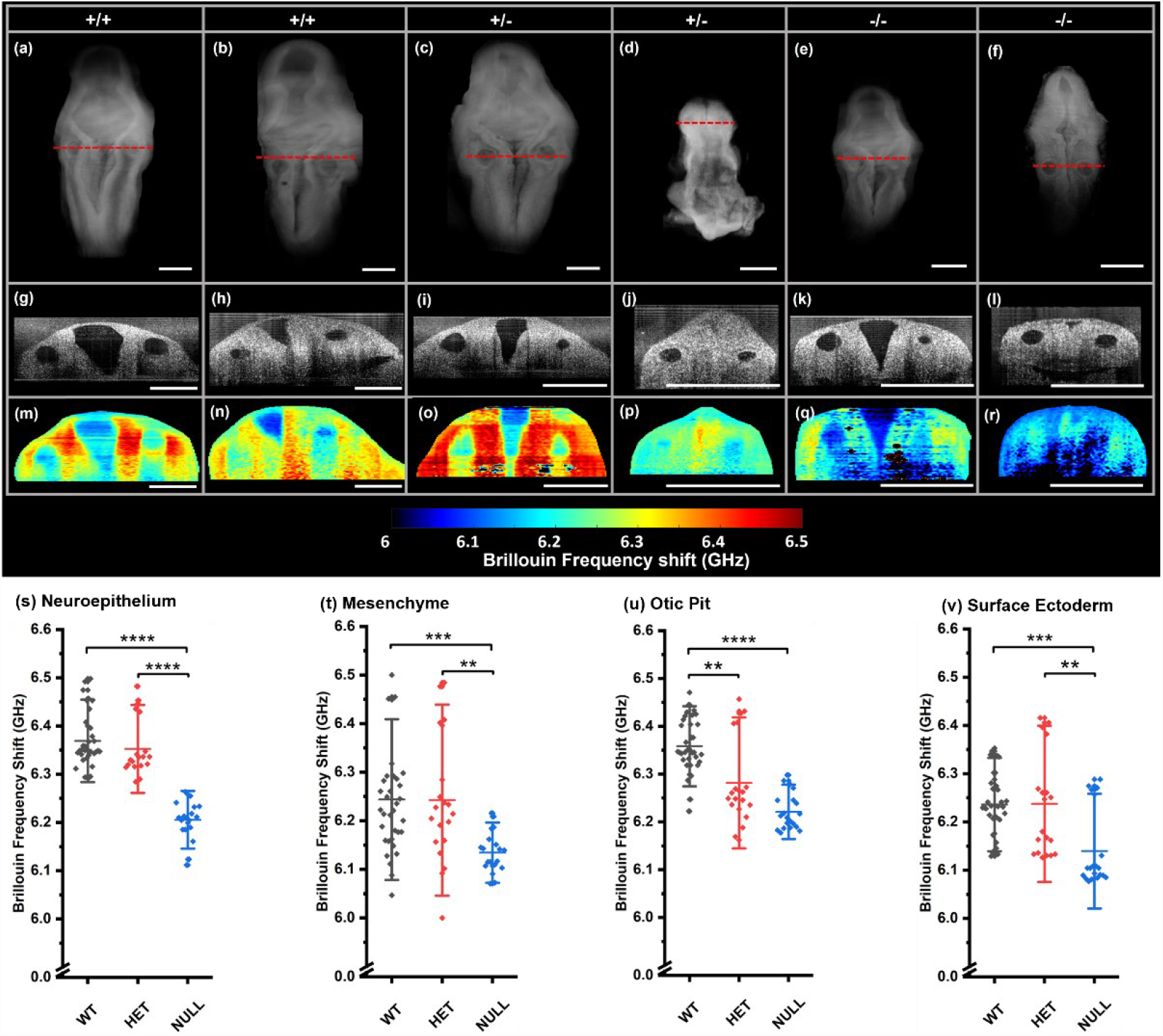
Mthfd1l ablation decreases tissue stiffness in E10.5 embryos. (a-f) 3D-OCT images showing the hindbrains of Mthfd1l embryos at E10.5. (g-l) 2D-OCT optical sections showing the neural folds at rhombomere 5 located near the otic pits. (m-r) Brillouin frequency shift images represent tissue stiffness in the anatomical areas identified by the corresponding 2D-OCT optical sections. Region-wise average Brillouin frequency shift of wild-type (WT), heterozygous (HET), and nullizygous (NULL) embryos at the (s) neural tube neuroepithelia, (t) adjacent paraxial mesenchyme, (u) otic pit, and (v) non-neural surface ectoderm region. *, **, ***, **** indicates P < 0.05, P < 0.01, P < 0.001, P <0.0001, respectively, by pair-wise Dunn’s test. Scale bar is 0.5 mm.

Early during embryonic development, it is expected that a relatively soft and flexible extracellular matrix exists, facilitating neural crest cell migration and a general rearrangement of the embryonic shape. As the embryo develops and the cells further differentiate, the cytoskeleton and the extracellular matrix become more organized, increasing tissue stiffness(*43*).

The embryos at E9.5 had an average Brillouin frequency shift of 6.18±0.06 GHz for the wild-type embryos (n=4), 6.14±0.06 GHz for heterozygous embryos (n=6), and 6.09±0.05 GHz for nullizygous embryos (n=6). The mice embryos at E10.5 had an average Brillouin frequency shift of 6.28±0.09 GHz for the wild-type embryos (n=6), while heterozygous embryos (n=5) had average shifts of 6.24±0.11 GHz, and nullizygous embryos (n=5) presented with shifts of 6.15±0.07 GHz (Figure 3, m-r). Considering the developmental window between E9.5 and E10.5 embryos, the Brillouin frequency shift of the developing neural tube was lower, as expected, in both heterozygous and nullizygous *Mthfd1l* mutant embryos when compared to the wild-type. Three-dimensional structural imaging with OCT at E9.5 showed normal NTC in the wild-type embryos (Figure 2, a-b), aberrant neural tube development in heterozygous (Figure 2, c-d), and a characteristic open or wavy neural tube in the nullizygous embryos (Figure 2, e-g). At E10.5, 3D structural imaging with OCT showed the characteristic growth delay expected in *Mthfd1l* mutants, including aberrant NTC in the mutants (Figure 3, e-f).

### Brillouin-OCT imaging of tissue stiffness rescued by formate supplementation

Mitochondrial loss of formate production is expected after *Mthfd1l* gene ablation. After determining that aberrant NTC in the *Mthfd1l* knockout mouse lineage is associated with alterations in tissue stiffness, we sought to determine if periconceptional maternal formate supplementation would improve tissue stiffness in the mutant embryos. Pregnant dams were given *ad libitum* access to water containing Calcium formate to achieve a calculated dose of 5,000 mg calcium formate·kg^−1^·d^−1^ (Methods).

At E9.5, the 3D-OCT structural images of formate-supplemented embryos showed normal NTC in all the embryos analyzed (Figure 4, a-f), regardless of genotype. As expected, the 3D-OCT structural images obtained from E10.5 formate-supplemented mutant embryos also showed normal NTC in all the embryos analyzed (Figure 5, a-f). At E9.5, formate-supplemented embryos had an average Brillouin frequency shift of 6.20±0.04 GHz in wild-type embryos (n=3), while heterozygous embryos (n=3) had a shift of 6.17±0.04 GHz, and the nullizygous mutants (n=3) had a shift of 6.17±0.04 GHz. Similarly, at E10.5, the average Brillouin frequency shift in the formate-supplemented wild-type embryos (n=3) was 6.25±0.07 GHz, in the heterozygous embryos (n=3) it was 6.25±0.06 GHz, and in the nullizygous mutant embryos (n=3) it was 6.21±0.06 GHz. Therefore, proper tissue stiffness as quantified by the Brillouin frequency shift was rescued in the *Mthfd1l* mutant embryos after formate supplementation (Figure 5, m-r).

**Figure 4.**
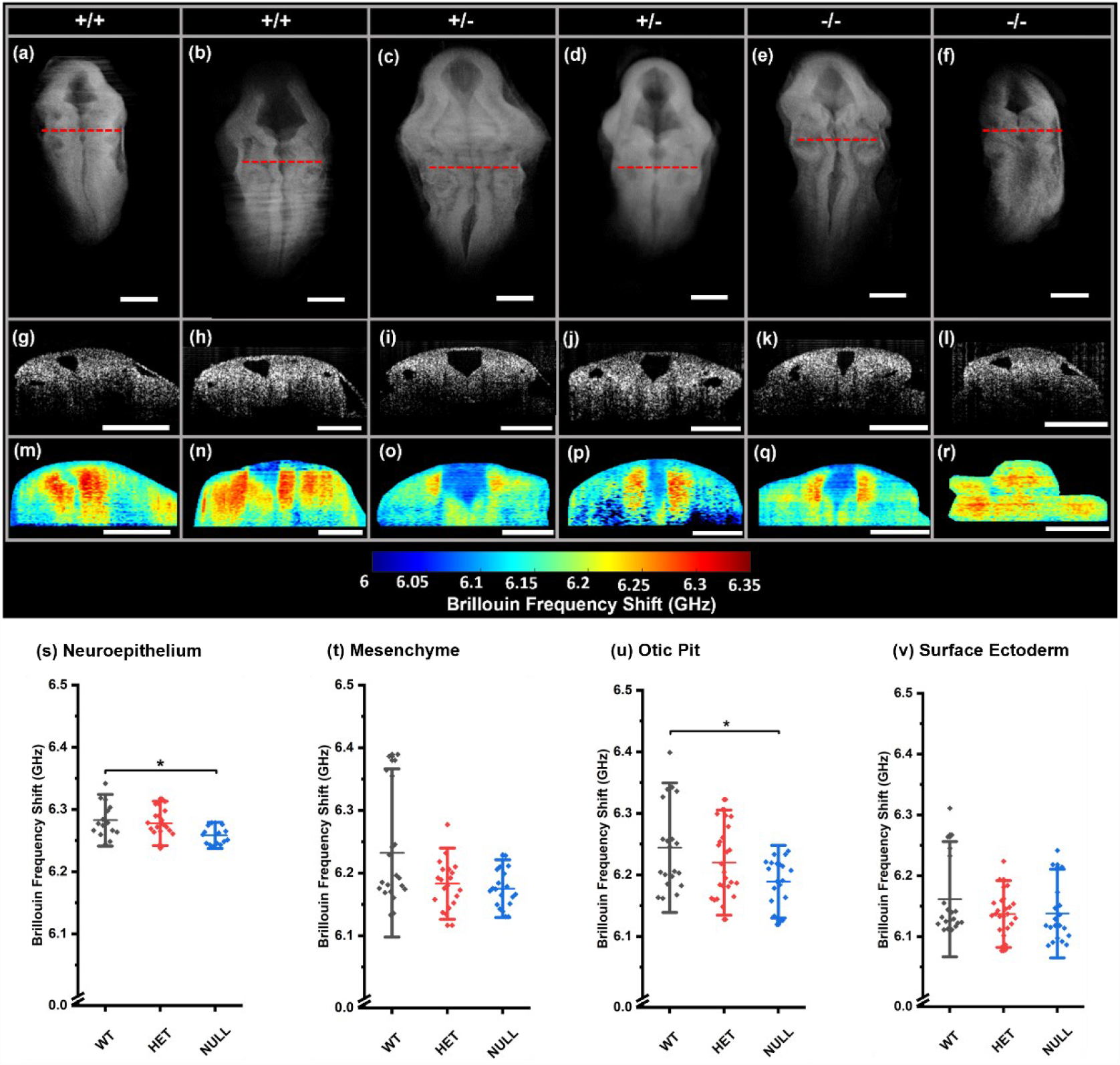
Formate supplementation improves tissue stiffness in E9.5 embryos. (a-f) 3D-OCT images showing the hindbrains of formate-supplemented Mthfd1l embryos at E9.5. (g-l) 2D-OCT optical sections showing the neural folds at rhombomere 5 located near the otic pits. (m-r) Brillouin frequency shift images represent tissue stiffness in the anatomical areas identified by the corresponding 2D-OCT optical sections. Region-wise average Brillouin frequency shift of wild-type (WT), heterozygous (HET), and nullizygous (NULL) embryos at the (s) neural tube neuroepithelia, (t) adjacent paraxial mesenchyme, (u) otic pit, and (v) non-neural surface ectoderm region. Formate supplementation in E9.5 embryos overcame the difference in the stiffness in all regions except for the neuroepithelium and otic pit region. * indicates P < 0.05 by pairwise Dunn’s test. The scale bar is 0.25 mm.

**Figure 5.**
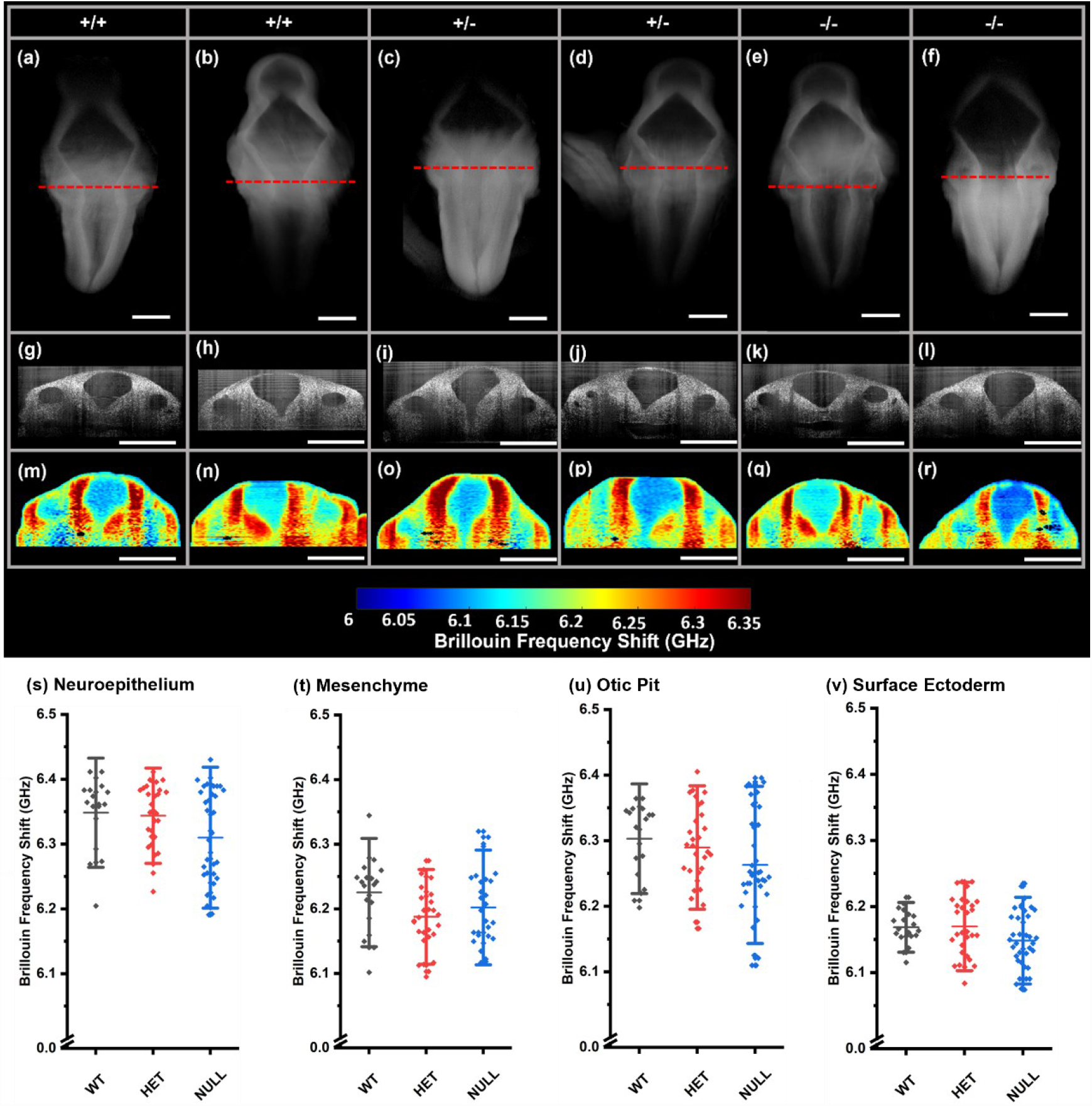
Formate supplementation improves tissue stiffness in E10.5 embryos. (a-f) 3D-OCT images showing the hindbrains of formate-supplemented Mthfd1l embryos at E10.5. (g-l) 2D-OCT optical sections showing the neural folds at rhombomere 5 located near the otic pits. (m-r) Brillouin frequency shift images represent tissue stiffness in the anatomical areas identified by the corresponding 2D-OCT optical sections. Region-wise average Brillouin frequency shift of wild-type (WT), heterozygous (HET), and nullizygous (NULL) embryos at the (s) neural tube neuroepithelia, (t) adjacent paraxial mesenchyme, (u) otic pit, and (v) non-neural surface ectoderm region. Formate supplementation in E10.5 embryos overcame the difference in stiffness in all the regions. The scale bar is 0.5 mm.

### Analysis of tissue stiffness during neural tube development in *Mthfd1l* embryos

The complete tissue stiffness analysis is provided in the supplementary information. For all *Mthfd1l* supplemented and non-supplemented embryos, the Brillouin frequency shift of the neural tube neuroepithelia, otic pit, non-neural surface ectoderm, and adjacent paraxial mesenchyme was statistically tested. The data were tested using Kruskal Wallis ANOVA’s and pairwise Dunn’s tests to see whether there was a significant difference between genotypes (Supplementary Table 1). At both the E9.5 and E10.5 stages for non-supplemented embryos, the average stiffness in the different regions of the wild-type embryo was greater in the wild-types and heterozygous embryos than in the nullizygous mutant embryos (Figure 2, v-y; Figure 3, s-v). The Brillouin frequency shift of all regions in similarly staged wild-type embryos was significantly different from that of nullizygous mutant embryos. Whereas, for E9.5 stage supplemented *Mthfd1l* embryos, the adjacent mesenchyme and surface ectoderm were not significantly different between wild-type, heterozygous, and nullizygous mutants showing that formate supplementation overcame the difference in stiffness in these regions. However, in these embryos, the neural tube neuroepithelia and the otic pit area of wild-type were still significantly different than the nullizygous embryos indicating that the supplementation at this stage did not overcome the stiffness difference completely in these regions (Figure 4, s-v). Furthermore, there was no significant difference in stiffness between wild-type, heterozygous, and nullizygous mutant embryos at stage E10.5, implying full stiffness recovery with formate supplementation (Figure 5, s-v) at this stage, as compared to E9.5. All plots shown in Figures 2-5 depict the mean and standard deviation of the Brillouin frequency shift.

### Immunohistochemical analysis of neural progenitors in *Mthfd1l* embryos

The importance of *Mthfd1l* gene expression during neural plate development and the elevated levels of *Mthfd1l* expression in the neuroepithelium, together with a variety of aberrant mutant phenotypes associated with decreased tissue stiffness during NTC(*13, 14*), led us to hypothesize that the population of neural progenitor cells would decrease following the overall growth delay and characteristic asymmetric neuroepithelium bulges observed in the wavy areas of the neural tube in mutant embryos. We anticipated that the neural fate from Pax6 positive progenitor cells would be disrupted along the embryonic antero-posterior axis. To test this hypothesis, we performed an evaluation of the neural progenitor fate using immunostaining against Pax6 and Tubb3 (neurofilament-Tuj1). Figure 6(a-c) shows that the population of Pax6 positive progenitor cells decreased in *Mthfd1l* mutant embryos, and consequently, the number of differentiated neurons along the body axis is significantly decreased. Our data therefore demonstrate that formate supplementation, shown in Figure 6(j-l) rescues not only the abnormal levels of Pax6 during neural tube development but also reestablishes neuronal differentiation from Pax6-positive progenitor cells in E9.5 embryos.

**Figure 6.**
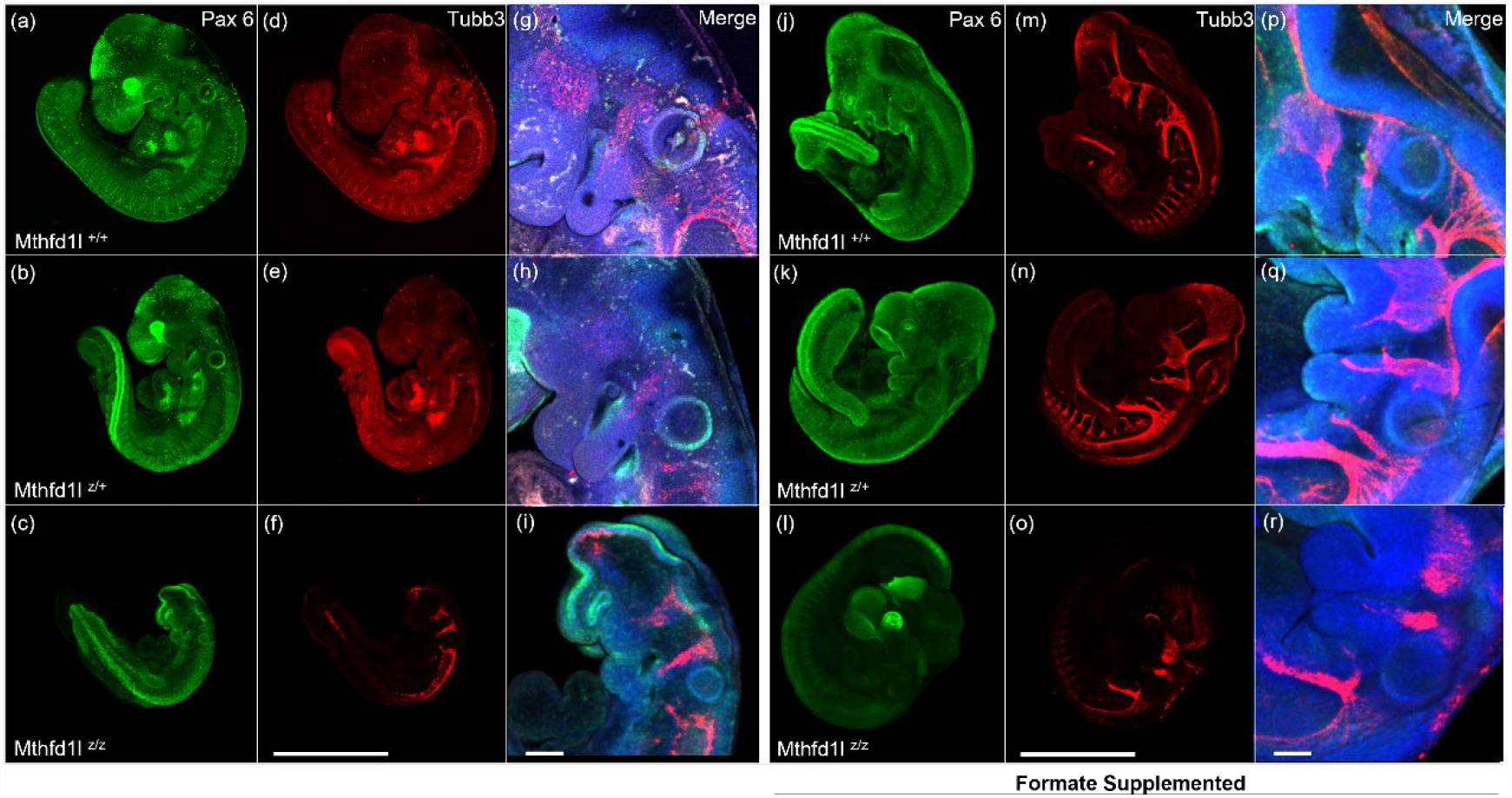
Mthfd1l ablation decreases neuronal differentiation in E9.5 embryos. Pax6-positive progenitor cells are decreased in Mthfd1l mutant embryos. Decreased neural differentiation is also detected along the body axis using Tubb3 antibody. Formate supplementations reestablish Pax6-positive progenitor cells and respective neural differentiation.

To determine whether the importance of mitochondrial formate was restricted to neuronal cell differentiation or if it could be correlated with tissue-specific differences in the *Mthfd1l* ablated mice, we also examined somitogenesis using Pax3 staining and the migration streams of Sox10 positive neural crest cells throughout the embryonic body plan. Pax3 immunostaining indicates that somitogenesis is abnormal in the *Mthfd1l* mutants. The immunostaining of Sox10 positive neural crest cells indicates decreased neural crest migration along the antero-posterior axis, but the general orientation of dorso-ventral migration streams did not change (Figure 7). Furthermore, we show that maternal supplementation with calcium formate rescues somitogenesis and Sox10-positive neural crest migration along the antero-posterior axis in embryos lacking a functional *Mthfd1l* gene (Figure 7).

**Figure 7.**
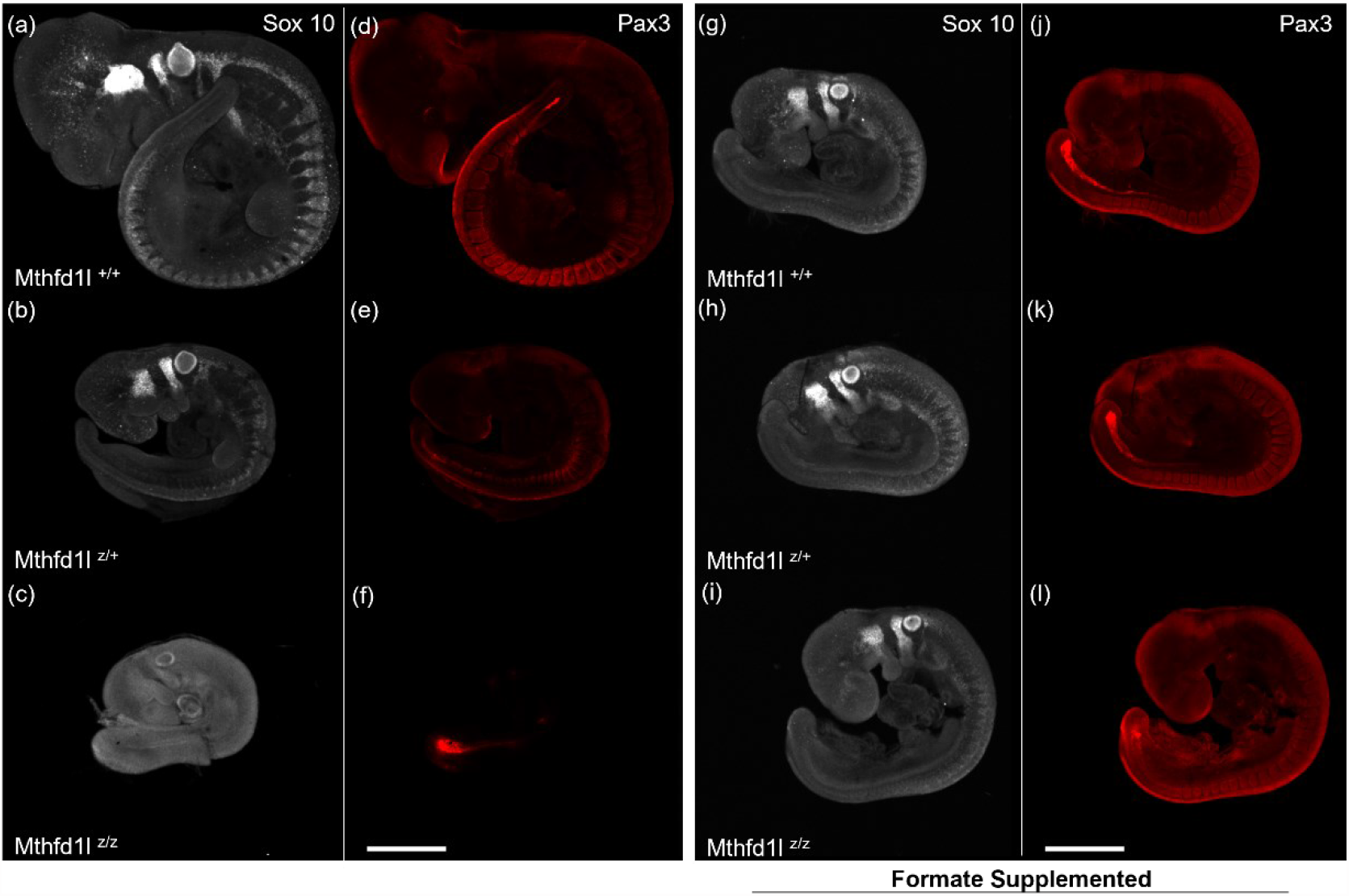
Mthfd1l ablation disrupts neural crest cell migration and somitogenesis in E9.5 embryos. Sox 10 positive neural crest cells are detected in decreased levels in Mthfd1l mutant embryos. Pax3 staining of somites indicates abnormal somitogenesis. The results should describe the experiments performed and the findings observed. The results section should be divided into subsections to delineate different experimental themes. Subheadings should be descriptive phrases. All data must be shown either in the main text or in the Supplementary Materials.

## Discussion

Synchronized development of the neural tube and adjacent anatomical structures is regulated by the composition of the extracellular environment, the activation of gene co-expression network modules, and the biomechanical forces arising from the cytoskeleton, cell-cell adhesions, and cell-matrix adhesions. These structures interact mechanically with one another to define an organized shape that is foundational for the development of an embryo. Understanding and imaging the biomechanical properties underlying embryo morphogenesis at the cellular or tissue level is a challenging problem due to a lack in developing instruments that could be adapted to measure the distribution of mechanical forces *in vivo*(*16, 44, 45*). In this study, we used a multimodal co-aligned OCT-Brillouin system(*42*) to measure and map tissue stiffness in *Mthfd1l* deficient mouse embryos with and without maternal formate supplementation at two distinct development stages. Structural data obtained using 3D-OCT imaging during hindbrain neurulation revealed genotype-dependent alterations in the distribution of tissue stiffness consistent with the abnormal NTC phenotypes observed in this mouse model (Figures 2 e-g and 3 d-f).

Following the original description by Momb and colleagues(*14*), our morphological analysis of mutant embryos using 3D-OCT provides a significantly more robust confirmation of the classic *Mthfd1l* wavy neural tube phenotype (Figures 2e, 3f), where the neural tube epithelia are compacted in an abnormal wavy line beginning at the cranial region and extending throughout the most caudal areas, disrupting normal NTC. We propose that the existence of these characteristic asymmetric bulges along the embryonic dorsal midline is due to a lack of neuroepithelial stiffness and alterations in stiffness of adjacent tissues, as well as failures located at the biomechanical network interlinking actomyosin contractility, turnover, or cytoskeletal assembly, which disrupts the connection of both neuroepithelial walls at their leading edge by the surrounding non-neural surface ectoderm(*46, 47*). Interestingly, a similar “wavy” neural tube phenotype (in some cases referred to as a “tortuous” or “kinked” neural tube or spine, has been observed in several other genetic mouse models of NTDs, including mutants of the chromatin regulators *Gmnn* and Fbxl10(*48-50*), a methyltransferase necessary for ribosome biogenesis, *Emg1*(*51*), and an apoptosis regulator, *Siva* (*52*). Notably, the phenotypes in each of these models are accompanied by increased apoptosis, decreased cellular proliferation, or altered differentiation of neural progenitors, similar to what is observed in *Mthfd1l* mutants. This information suggests some common mechanisms indicating that cellular density and composition of tissues influence their stiffness during biomechanical processes like NTC.

In this study, we used a novel imaging technique utilizing high-resolution 2D-OCT as a starting point to capture Brillouin-shifted scattered light at transversal sections crossing a segment of the embryonic hindbrain. Differences in the Brillouin frequency shift detected in these sections identified the neural tube epithelial walls, surface non-neural ectoderm, otic pits, and adjacent mesenchyme. Analysis of the Brillouin frequency shift at E9.5 showed that the surface ectoderm already connected the neural folds at the level of the rhombomere 5 in all the embryos analyzed, except for the nullizygous embryos represented in Figure 2 (o-u). Tissue stiffness analysis at the neuroepithelial walls, otic pit, non-neural surface ectoderm, and adjacent paraxial mesenchyme showed a significative loss of stiffness in the *Mthfd1l* mutants at E9.5 and E10.5 (Figures 2 and 3). Proper NTC requires an initial contact established between the apposed neural fold tips, which is only possible via cell protrusions and membrane ruffles, which will connect the pseudostratified neuroepithelium and the squamous surface ectoderm. The surface ectoderm is also responsible for building a structural backbone made of high-tension actomyosin cables to pull the leading edge of both neuroepithelial walls together across the dorsal midline(*53, 54*). Surprisingly, stiffness at the surface ectoderm is lower in *Mthfd1l* mutants at E9.5 and E10.5, indicating that the opposed neuroepithelial walls cannot present resistance against the biomechanical forces generated by the surface ectoderm actomyosin backbone (Figures 2 and 3).

The Brillouin frequency shift captured from wild-type embryos at E9.5 and E10.5 clearly shows that the otic pit and the neural tube epithelial layers are stiffer than the adjacent mesenchyme and surface ectoderm (Figures 2 and 3). The mesenchyme at the rhombomere 5 transverse section receives a higher contribution of neuroblasts, which are delaminated from the otic pit epithelia, and a much smaller contribution of cells from the neural crest, which may still contribute to inner ear development in small amounts(*55, 56*). At E9.5 and E10.5, the otic pits and the adjacent mesenchyme of mutant embryos were less stiff than the wild-type. As shown in Figures 4 and 5, formate supplementation also reestablishes stiffness in the otic pits as well as the adjacent mesenchyme, indicating that these tissues contribute to the balance of stiffness during the development of posterior hindbrain structures.

Shin and colleagues reported in 2019 that deletion of *Mthfd1l* causes reduced cranial mesenchyme density at E8.5, which is a stage prior to hindbrain neuropore formation and subsequent hindbrain neuropore closure(*32*). Tissue stiffness measured at this scale (tens of micrometers) is heavily influenced by cellular density(*57*). Although Brillouin microscopy measurements depend on other factors like hydration, it is still capable of sensing tissue stiffness based on other factors(*42, 57*). Earlier results in wild-type embryos have shown a clear difference between the cell-dense neuroepithelial region (higher Brillouin frequency shift) and the less cell-dense mesoderm layer (lower Brillouin frequency shift)(*42, 58*). Our results explicitly show similar observations where the cell-dense neuroepithelial layers had a greater Brillouin shift compared to the other parts of the embryo. In the mutant embryos, there was a reduction in the Brillouin shift in the neuroepithelial region as well as the mesenchyme layer, indicating a lower stiffness than in the wild-type embryos. Embryos that were supplemented with formate had greater Brillouin frequency shifts in the mutant than non-supplemented embryos, indicating improvement in the stiffness of the neural tube tissue with supplementation along with a corresponding decrease in the occurrence of NTDs. Therefore, the tissue needs to have sufficient stiffness for proper NTC.

The decreased Brillouin shift measured in the adjacent mesenchyme at E9.5 and E10.5 (Figures 4 and 5) supports a scenario where the extracellular matrix and cytoskeletal components derived from mesenchymal cells are softer than the wild-type, probably due to weaknesses in the neuroepithelial walls, which are unable to support the biomechanical forces coming from the actomyosin backbone established by the surface ectoderm. This imbalance of forces established between mutant neuroepithelial walls, the surface ectoderm, and the adjacent mesenchyme may explain the appearance of the characteristic neural tube wavy phenotype in mutants of the *Mthfd1l* lineage (Figure 2e).

Collectively, considering the developmental window between E9.5 and E10.5 embryos, the Brillouin frequency shift of the developing neural tube was restored after formate supplementation in both heterozygous and nullizygous *Mthfd1l* mutant embryos, as expected (Figure 5). Decreased stiffness levels were detected specifically at the neural epithelial walls, which can be correlated with the importance of *Mthfd1l* for proper neural plate development. Previous work in this mouse lineage indicated that the *Mthfd1l* gene is highly expressed in the mouse embryonic neuroepithelium(*13, 14*). Considering the aberrant neural tube phenotypes associated with a significant decrease in the neuroepithelial stiffness during NTC in *Mthfd1l* mutants, we decided to validate the connection between stiffness and cell fate aberrations in *Mthfd1l* knockout neural progenitor cells. We chose immunostaining against Pax6 and Tubb3 (Tuj1 monoclonal serum against neurofilaments). Pax6 and Tubb3 provide information about neuronal cells and their differentiation and development of neurofilaments. Figure 6 indicates that neuronal differentiation was disrupted at the cranial ganglia derived from rhombomeres 2, 4, 6, 7, and 8, and along the embryonic posterior axis in the absence of mitochondrial formate. Lower neural differentiation was also detected along the body axis using the Tubb3 antibody, as shown in Figure 6. Both the Pax6 and Tubb3 results corroborate with the lower stiffness measurements in the *Mthfd1l* mutants. Neuronal differentiation from Pax6 positive progenitor cells was reestablished along the entire body axis after formate supplementation of *Mthfd1l* mutants (Figure 6), also validating the improved stiffness observed in Brillouin results.

It is well known that cranial neural crest cell migration happens at the even rhombomeres 2 and 4 anteriorly located in the mouse hindbrain, avoiding the constricted areas in the odd rhombomeres 1, 3, and 5(*59, 60*). The otic pit is an embryonic structure located adjacent to rhombomere 5, a hindbrain area where the mesenchyme does not receive a significative contribution of the neural crest population of cells(*55*), but instead is infiltrated by neuroblast cells delaminated from the otic pit(*56*). Considering the decreased tissue stiffness detected in the mesenchyme of *Mthfd1l* embryonic mutants, we prepared immunostaining experiments to check for alterations in the migration streams of Sox10 positive neural crest cells throughout the rhombomeres 2 and 4 and along the embryonic antero-posterior body plan. Sox10 positive neural crest migration is disrupted in the absence of *Mthfd1l*, but the general orientation of dorso-ventral migration streams did not change. Shin and colleagues also reported that deletion of *Mthfd1l* does not affect neural crest cell specification, using immunostaining against Sox9 at E8.5. The SRY-Box transcription factor 9 (Sox9) is a broader neural progenitor marker, and its expression pattern is not restricted to the neural crest population. Sox9 is expressed all along the developing neural tube axis and activates genes that will induce the migration of neural crest cells(*32, 61*). Sox10 positive neural crest cell migration was also rescued by formate supplementation (Figure 7).

Considering the importance of *Mthfd1l* for proper neural differentiation, we also examined somitogenesis to explore a hypothesis where the importance of mitochondrial formate could be correlated with tissue-specific differences in the *Mthfd1l* ablated mice. Pax3 immunostaining indicates that somite segmentation is altered at E9.5, following the overall developmental delay observed in *Mthfd1l* mutants. Normal somite segmentation could be observed along the embryonic body plan after maternal formate supplementation (Figure 7).

During these experiments, we were able to characterize the biomechanical properties of four individual tissue compartments during hindbrain neuropore closure in mouse embryos lacking *Mthfd1l*. These data suggest that mitochondrial one-carbon metabolism is functionally critical to modulating embryonic tissue stiffness required for NTC. There are several reasons why this may be the case. Mitochondrial formate is essential for the synthesis of thymidine for the replication of DNA during cellular division. It has also been demonstrated in mouse embryonic stem cells that up to 75% of carbon units entering the methylation cycle are derived from mitochondrial formate(*13*), and as mentioned earlier, transgenic mouse lines with disruptions in certain demethylases or methyltransferases present with similar neural tube phenotypes as *Mthfd1l* knockouts. Additionally, mitochondrial one-carbon metabolism has been previously shown in other contexts to mediate redox homeostasis and promote cell survival. Finally, we demonstrate here that certain cell types do not properly differentiate in the absence of *Mthfd1l*. Thus, impairment of mitochondrial one-carbon metabolism and resulting formate production may alter the composition and density of embryonic tissues through a variety of mechanisms, ultimately disrupting their biomechanical properties. Our results show, for the first time, the distribution of tissue stiffness during this process and that the absence of formate significantly hinders neurulation, resulting in significant defects. Moreover, supplementation of formate dramatically reduced any stiffness and phenotypic abnormalities, further enforcing the importance of formate.

## Materials and Methods

### Animal Husbandry

All mice were maintained on the C57BL/6 background and housed in a 16-hour light:8-hour dark-light cycle. All animal experiments were conducted in accordance with Institutional Animal Care and Use Committees approved protocols at the Baylor College of Medicine and the University of Houston. *Mthfd1l*, an essential enzyme for tetrahydrofolate synthesis inside the mitochondria was ablated in this lineage(*13, 14*). Male and female heterozygous mice were mated overnight, and the presence of the vaginal plug was checked every morning. The morning, when the vaginal plug was observed, was considered as E0.5. The pregnant mice were euthanized at E9.5 and E10.5, embryos were dissected, and the embryos were kept in 100% rat serum at Baylor College of Medicine. The fresh embryos were then transferred to the University of Houston within 30 minutes. The yolk sac around the embryos was removed, and the embryos were mounted on a 1.5% agarose plate filled with the culture media. The embryos were aligned with their neural tube side up to acquire Brillouin-OCT measurements.

### Multimodal Brillouin-OCT system

The multimodal Brillouin-OCT system has been described in a previous article(*42*). Briefly, the home-built system consists of swept source OCT sub-system, a Brillouin microscopy sub-system with a dual virtually imaged phase array (VIPA) spectrometer, and a combined scanning arm. The Brillouin sub-system utilized a 660 nm single-mode laser source. The incident power on the sample was 35 mW. The collected backscattered light from the sample was transferred to the dual VIPA spectrometer, and an electron multiplying charge coupled device camera was used to detect the Brillouin frequency shift of the sample. The camera acquisition time was 0.2 s. The system was calibrated with standard materials such as water, acetone, and methanol. The sample was imaged with an achromatic doublet with 0.25 NA, resulting in an axial resolution of ∼36 μm and lateral resolution of ∼3.8 μm. The swept source OCT sub-system had a central wavelength of ∼1310 nm, a scan rate of 50 kHz, a scan range of ∼105 nm, and ∼8 mW incident power on the sample. The lateral and axial resolutions were ∼17.5 μm and ∼10 μm in air, respectively. Light from both systems was combined using a dichroic mirror, and galvanometer-mounted mirrors scanned the beam across the sample. For Brillouin imaging, the sample was stepped by a motorized vertical stage. A custom instrumentation software was developed that utilizes the OCT structural image to guide Brillouin imaging(*42*).

### Embryo genotyping

The embryo yolk sac tissues were used to perform genotyping to differentiate the embryos as wild-type (*Mthfd1l*^+/+^), heterozygous (*Mthfd1l*^+/-^), and nullizygous mutant (*Mthfd1l*^-/-^). After imaging with the Brillouin-OCT system, the embryos were fixed in 4% paraformaldehyde solution and transferred for further imaging.

### Formate supplementation

Calcium formate at 200 μM concentration was added to the drinking water of the female *Mthfd1l*^+/-^ mice for 10 days prior to the first attempt at mating. Considering an average intake of 5 mL water per day for an average of 25 g mouse, this dose delivers around 1300 mg of formate. After 10 days of supplementation, these mice were set up for timed mating, and at E9.5 and E10.5 the embryos were imaged using the Brillouin-OCT system.

### Immunohistochemistry and DAPI Staining

Pax6 (42-6600, ThermoFisher) and Tubb3 (2H3-Tuj1, DSHB); Sox10 and Pax3 (DSHB) whole-mount immunostaining was performed on E9.5 embryos, after fixation in 4% PFA and dehydration to 100% methanol. Endogenous peroxidase activity was blocked by incubation in Dent’s bleach for 1 hr at room temperature. Embryos were rehydrated to TBS containing 0.1% Tween-20 (TBST) and incubated in a 1:500 dilution of anti-Pax6 and anti-Tuj1 overnight at room temperature. Embryos were washed five times in PBST, and then incubated with a 1:1000 fluorescent secondary antibodies. For general cell visualization, the embryos were incubated overnight in DAPI (4′,6-diamidine-2-phenylidole-dihydrochloride; 2 μg/ml) to label all the nuclei. Embryos were cleared in 25% glycerol and imaged using a Nikon CSU-W1 Yokogawa spinning disc confocal microscope. The obtained Z-stacks were projected at maximum intensity and exported as TIFF files(*8*).

## Author contributions

Y.S.A., Conceptualization, Formal Analysis, Investigation, Methodology, Visualization, Writing – original draft, Review, and Editing

C.D.C., Conceptualization, Formal Analysis, Investigation, Methodology, Resources, Visualization, Writing, Review, and Editing

M.S., Formal Analysis, Methodology, Visualization, Review, and Editing

A.W.S., Formal Analysis, Methodology, Visualization

J.S., Methodology, Visualization, Review, and Editing

S.R.A., Supervision, Formal Analysis, Methodology, Visualization, Review, and Editing

G.S., Supervision, Writing – review and editing, Project administration, Funding acquisition

R.H.F., Supervision, Writing – review and editing, Project administration, Funding acquisition K.V.L., Supervision, Writing – review and editing, Project administration, Funding acquisition

## Data and materials availability

All data, code, and materials used in the analyses is available upon reasonable request.

