## Supplemental Figures for "Aberrant tissue stiffness impairs neural tube development in Mthfd1l mutant mouse embryos"

### Supplementary Materials

The work described in this study demonstrates a novel, multimodal imaging technique combining optical coherence tomography (OCT) and Brillouin light scattering microscopy to evaluate the structural and biomechanical changes of the neural tube in a murine neural tube defect (NTD) model. Hence, we developed a co-aligned Brillouin-OCT system with customized instrumentation software *(42)*. We adapted, optimized, and combined the two technologies to build a single Brillouin-OCT instrument for structurally guided tissue stiffness mapping of neurulation in Mthfd1l knockout mouse embryos with high resolution. The schematic of multimodal Brillouin-OCT is shown in Figure 1-a (ADC - analog to digital convertor; BPD - balanced photodetector; Br - Brillouin; C, C1, C2 - collimator; CL - collection lens; DAC - digital to analog converter; DM - dichroic mirror; EMCCD - electron multiplying charged coupled device camera; FC - fiber coupler; GS - galvo scanning mirrors; λ/4 - quarter-wave plate; OL - objective lens; PBS - polarizing beam splitter; PC - polarization controller; RM - reference mirror; SPEC - dual-stage virtual image phase array spectrometer; VA - variable attenuator).

<u>Area selection for tissue stiffness analysis during neural tube development in Mthfd1l embryos</u>

It is critical to know which anatomical region is being imaged with Brillouin microscopy. Hence, OCT was combined with Brillouin microscopy. Figure 1-b represents the flowchart of the imaging procedure to select the region of interest (ROI) for Brillouin imaging based on the OCT structural image. First, 3D-OCT images of a mouse embryo at a given embryonic stage were acquired (Figure 1-c). For all embryos, a 2D-OCT image of a coronal plane across the neural tube and otic pits region was acquired. Using the home-made instrumentation software, an ROI was chosen on the

2D-OCT image once a sufficient coronal plane was acquired, and this ROI was utilized to obtain the Brillouin frequency shift map (Figure 1-d). Using linear interpolation, the Brillouin frequency shift image was rescaled to match the 2D-OCT image (Figure 1-f). The OCT image was normalized to [0,1], where 0 was the noise floor, and 1 was the 75th percentile of the intensities, and this image acted as a mask to select the pixels of the Brillouin frequency shift map (Figure 1-e). The average Brillouin frequency shift after masking was used for further analysis. Furthermore, the regional Brillouin frequency shift at the neural tube neuroepithelia, otic pit, non-neural surface ectoderm, and adjacent paraxial mesenchyme of all the embryos in Mthfd1l supplemented and the non-supplemented category was selected as shown in Figure 1(h).

Statistical analysis of tissue stiffness during neural tube development in Mthfd1l embryos

Region-wise tissue stiffness analysis

The region-wise Brillouin frequency shift at the neural tube neuroepithelia, otic pit, non-neural surface ectoderm, and adjacent paraxial mesenchyme of formate-supplemented and non-supplemented Mthfd1l embryos were analyzed using Kruskal Wallis ANOVA. The Brillouin frequency shifts of neuroepithelia for E9.5 non-supplemented Mthfd1l embryos were significantly different as a function of genotype using Kruskal Wallis ANOVA ($\chi^2$=75.667, P<<0.001). A similar difference was observed for the neuroepithelia of E10.5 stage non-supplemented Mthfd1l embryos ($\chi^2$= 45.524, P<<0.001). The Brillouin frequency shifts of adjacent mesenchyme for age-matched non-supplemented Mthfd1l embryos were significantly different as a function of genotype using Kruskal Wallis ANOVA ($\chi^2$=43.716, P<<0.001 for E9.5; $\chi^2$=18.654, P<<0.001 for E10.5). Similarly, the Brillouin frequency shifts of the otic pit for age-matched non-supplemented Mthfd1l embryos were significantly different as a function of genotype using Kruskal Wallis ANOVA ($\chi^2$=33.935, P<<0.001 for E9.5; $\chi^2$=35.656, P<<0.001 for E10.5). Moreover, the Brillouin frequency shifts of surface ectoderm for age-matched non-supplemented Mthfd1l embryos were significantly different as a function of genotype using Kruskal Wallis ANOVA ($\chi^2$=48.061, P<<0.001 for E9.5; $\chi^2$=17.851, P<0.001 for E10.5). The pairwise difference among the age matched Mthfd1l genotypes was tested by Dunn's test, and the results are shown in Table 1. The Brillouin frequency shifts of neuroepithelia for formate-supplemented E9.5 Mthfd1l embryos were different as a function of genotype using Kruskal Wallis ANOVA ($\chi^2$=8.193, P=0.017), and a pairwise difference was observed between the wild-type and nullizygous mutants (Table 1). The Brillouin frequency shifts of adjacent mesenchyme, otic pit, and surface ectoderm for formate-supplemented E9.5 Mthfd1l embryos were not different as a function of genotype using Kruskal Wallis ANOVA ($\chi^2$=4.467, P=0.107; $\chi^2$=5.814, P=0.055; $\chi^2$=1.632, P=0.442), and a pairwise analysis by Dunn's Test is shown in Table 1. The Brillouin frequency shifts of neuroepithelia, adjacent mesenchyme, otic pit, and surface ectoderm for E10.5 supplemented Mthfd1l embryos were not different as a function of genotype using Kruskal Wallis ANOVA ($\chi^2$=4.959, P=0.084; $\chi^2$=4.817, P=0.090; $\chi^2$=3.812, P=0.149; $\chi^2$=5.681, P=0.058 respectively). Region-wise tissue stiffness analysis is more precise to see the changes in the tissue stiffness before and after supplementation in Mthfd1l knockout embryos. Moreover, this analysis showed that the supplemented embryos recovered stiffness in different regions of embryos at the E10.5 stage. Embryos that were supplemented with formate had greater Brillouin frequency shifts in the mutant than non-supplemented embryos, indicating improvement in the stiffness of the neural tube tissue with supplementation along with a corresponding decrease in the occurrence of NTDs. Therefore, a specific degree of stiffness is necessary for proper neural tube closure. Formate supplementation at E10.5 stage embryos showed significant improvement in the stiffness in nullizygous embryos for all the regions compared to E9.5 stage supplemented embryos, indicating the effect of supplementation is more visible in the later stages of embryos (Table 1). It will be interesting to explore tissue stiffness in supplemented embryos at a later developmental stage in future studies.

The whole field of view tissue stiffness analysis

Here, the average tissue stiffness for all embryos over the entire imaged field of view was calculated, as shown in Figure 1(c-g). Figure 2 plots the average Brillouin frequency shift of the individual groups of embryos separated by developmental stage, supplementation, and genotype. Statistical analysis using a Mann-Whitney U-test between formate-supplemented and non-supplemented Mthfd1l mouse embryos at E9.5 and E10.5 for each genotype is shown in Table 2. There was no significant difference in the average Brillouin frequency shift of the wild-type embryos with and without formate supplementation at E9.5. Similarly, there was no significant difference in the average Brillouin frequency shift of the heterozygous embryos with and without formate supplementation at E9.5. Similar observations were made for wild-type and heterozygous embryos at E10.5. However, at E9.5 and E10.5, the nullizygous mutant data showed a significant difference in the average Brillouin frequency shift of embryos that were supplemented with formate and those without supplementation, indicating a significant improvement in the neural tube stiffness in the nullizygous embryos, which was correlated with the presence of NTDs. This study underlines the role of measuring mechanical properties in the normal development of neural tube.

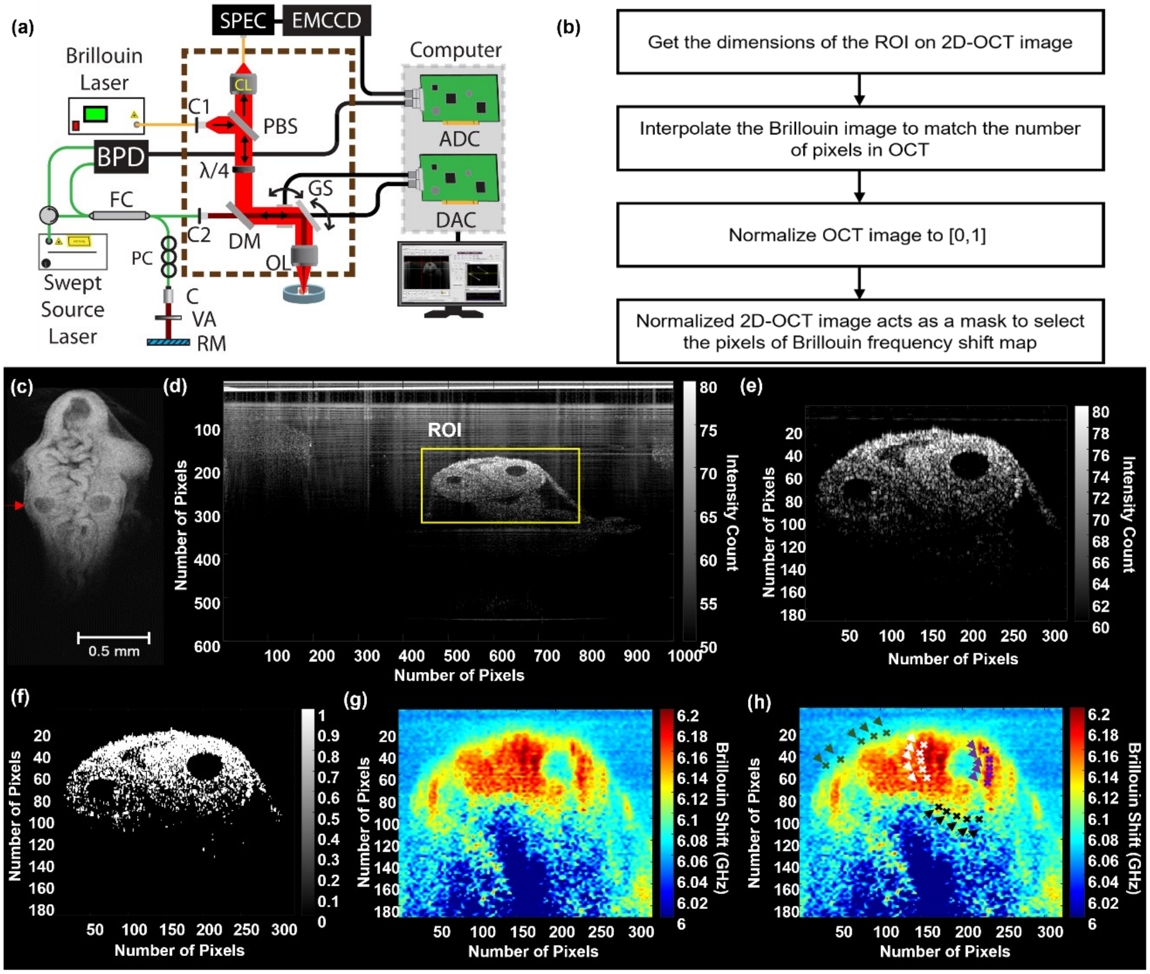


**Figure S1.** (a) Schematic of multimodal Brillouin-OCT system. (b) Flowchart showing steps to select the Brillouin frequency shift maps using OCT image. (c) 3D-OCT image of a Mthfd1l mouse embryo (d) 2D-OCT image acquired at red arrow shown in 3D-OCT image and yellow square indicated region of interest (ROI) selected at neural tube area. (e) 2D-OCT from the ROI. (f) Normalized 2D-OCT image. (g) Interpolated Brillouin frequency shift map to match the number of pixels in OCT. The dots on the Brillouin frequency shift map represents different regions: neuroepithelium (white), mesenchyme (black), otic pit (purple), surface ectoderm (green).

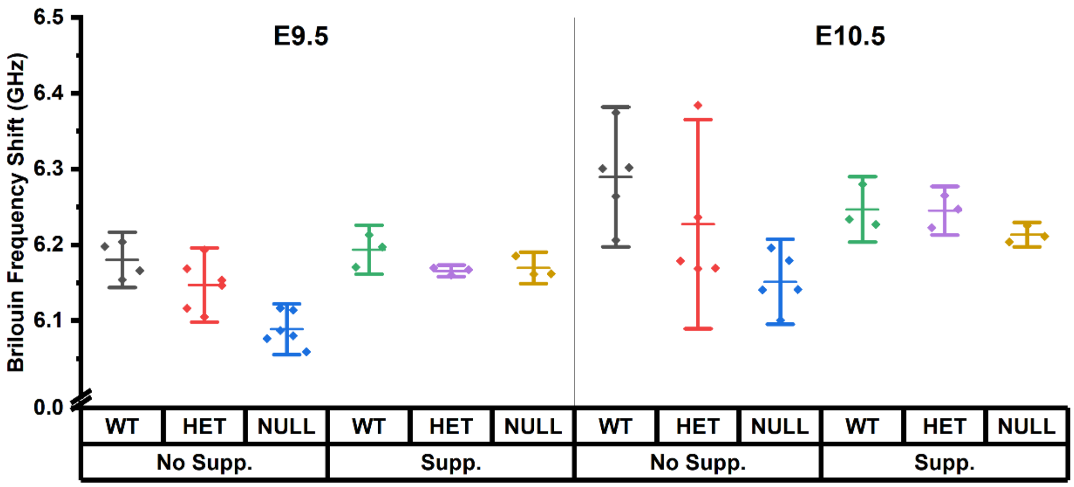


**Figure S2.** Average Brillouin frequency shift of wild-type (WT), heterozygous (HET), and nullizygous (NULL) Mthfd1l embryos without supplementation (No Supp.) and the formate-supplemented (Supp.) at E9.5 and E10.5.

**Table S1.** Pairwise statistical testing by Dunn's test (* denotes significant difference)

| | | Test Pair | Neuroepithelium | Mesenchyme | Otic Pit | Surface Ectoderm |
|---|---|---|---|---|---|---|
| E9.5 | Non-Supplemented Mthfd1l | WT-HET | **P=0.015*** | P=0.099 | P=0.229 | **P=0.014*** |
| | | WT-NULL | **P<0.001*** | **P<0.001*** | **P<0.001*** | **P<0.001*** |
| | | HET-NULL | **P<0.001*** | **P<0.001*** | **P<0.001*** | **P<0.001*** |
| | Supplemented Mthfd1l | WT-HET | P=1 | P=0.494 | P=0.798 | P=1 |
| | | WT-NULL | **P=0.026*** | P=0.114 | **P=0.049*** | P=0.606 |
| | | HET-NULL | P=0.054 | P=1 | P=0.465 | P=1 |
| E10.5 | Non-Supplemented Mthfd1l | WT-HET | P=0.444 | P=1 | **P=0.004*** | P=1 |
| | | WT-NULL | **P<0.001*** | **P<0.001*** | **P<0.001*** | **P<0.001*** |
| | | HET-NULL | **P<0.001*** | **P=0.002*** | P=0.061 | **P=0.003*** |
| | Supplemented Mthfd1l | WT-HET | P=1 | P=0.090 | P=1 | P=1 |
| | | WT-NULL | P=0.171 | P=0.351 | P=0.227 | P=0.194 |
| | | HET-NULL | P=0.221 | P=1 | P=0.466 | P=0.109 |

**Table S2.** Statistical analysis using Mann-Whitney U-test between formate-supplemented and non-supplemented Mthfd1l mouse embryos at E9.5 and E10.5 (* denotes significant difference)

| | | **Genotype** | | |
|---|---|---|---|---|
| | | WT | HET | NULL |
| **Stage** | E9.5 | p = 0.596 | p = 0.366 | **p = 0.028*** |
| | E10.5 | p = 0.371 | p = 0.371 | **p = 0.036*** |
